# Reconstructing Computational Dynamics from Neural Measurements with Recurrent Neural Networks

**DOI:** 10.1101/2022.10.31.514408

**Authors:** Daniel Durstewitz, Georgia Koppe, Max Ingo Thurm

## Abstract

Mechanistic and computational models in neuroscience usually take the form of systems of differential or time-recursive equations. The spatio-temporal behavior of such systems is the subject of dynamical systems theory (DST). DST provides a powerful mathematical toolbox for describing and analyzing neurobiological processes at any level, from molecules to behavior, and has been a mainstay of computational neuroscience for decades. Recently, recurrent neural networks (RNNs) became a popular machine learning tool for studying the nonlinear dynamics underlying neural or behavioral observations. By training RNNs on the same behavioral tasks as employed for animal subjects and dissecting their inner workings, insights and hypotheses about the neuro-computational underpinnings of behavior could be generated. Alternatively, RNNs may be trained *directly* on the physiological and behavioral time series at hand. Ideally, the once trained RNN would then be able to generate data with the *same temporal and geometrical properties* as those observed. This is called *dynamical systems reconstruction*, a burgeoning field in machine learning and nonlinear dynamics. Through this more powerful approach the trained RNN becomes a *surrogate* for the experimentally probed system, as far as its dynamical and computational properties are concerned. The trained system can then be systematically analyzed, probed and simulated. Here we will review this highly exciting and rapidly expanding field, including recent trends in machine learning that may as yet be less well known in neuroscience. We will also discuss important validation tests, caveats, and requirements of RNN-based dynamical systems reconstruction. Concepts and applications will be illustrated with various examples from neuroscience.

## Introduction

A long-standing tenet in theoretical neuroscience is that computations in the nervous system can be described and understood in terms of the underlying nonlinear system dynamics (Amit & Brunel, 1997; Brody & Hopfield, 2003; Brunel, 2000; Durstewitz, 2003; Durstewitz et al., 1999, 2000, 2021; Hodgkin & Huxley, 1952; Hopfield, 1982; Izhikevich, 2007; Machens et al., 2005; Miller, 2016; Rinzel & Ermentrout, 1998; Wang, 1999, 2002; Wilson, 1999; Wilson & Cowan, 1972). Related ideas may be traced back to work by McCulloch & Pitts (1943), Alan Turing (1948) and Norbert Wiener (1948) in the forties, and gained momentum in the early eighties through the seminal work by John Hopfield (1982) on embedding of memory patterns as fixed point attractors in simple recurrent neural networks. The beauty of Hopfield networks was that they delivered many properties of biological cognitive systems for free, such as automatic pattern completion, content-addressable memory retrieval through partial cues, or robustness to partial lesions and noise. Viewing neural computations through the lens of dynamical systems theory (DST) is particularly powerful since, on the one hand side, many (if not most) physical and biological processes are naturally formalized in terms of differential or difference equations (aka, dynamical systems [DS]). On the other hand, DS are computationally universal in the sense that they can mimic the operations of any computer algorithm (more technically, they are Turing complete: any Turing machine can be implemented as a DS (Branicky, 1995; Koiran et al., 1994)). Hence, DST offers both, a natural and mighty mathematical language for biochemical (Gutfreund, 1995), molecular (Bhalla & Iyengar, 1999, 2001; Leimkuhler & Matthews, 2015) and physiological (Durstewitz & Gabriel, 2007; Durstewitz & Seamans, 2002; Mackey & Glass, 1977; Rinzel & Ermentrout, 1998; Sherman, 2011) processes in their own right, as well as for understanding information processing and computation. It therefore provides a bridge between these levels of nervous system description.

Although the value of DST for explaining physiological and computational processes in the nervous system has been appreciated for a long time, until recently it was difficult to assess DS properties from neural recordings directly. Mostly, theory in neuroscience proceeded by first formulating mathematical models based on first principles (e.g., Hodgkin-Huxley kinetics; Hodgkin & Huxley, 1952) or their abstractions (e.g., FitzHugh-Nagumo equations; FitzHugh, 1955; Nagumo et al., 1962), which are then simulated and analyzed to yield insights and experimental predictions. The analysis of these often complex models can be very tedious in its own right. The prospects for applying DST tools in neuroscience have changed dramatically recently, however, with the advance of massive neural recording techniques on the one hand (Machado et al., 2022; Paulk et al., 2022; Steinmetz et al., 2021; Urai et al., 2022; Vogt, 2019), and powerful machine learning tools on the other (Brunton et al., 2016; Champion et al., 2019; Durstewitz, 2017b; Hernandez et al., 2020; Kass et al., 2014; Kim et al., 2021; Koppe et al., 2019; Kramer et al., 2022; Pandarinath et al., 2018; Paninski & Cunningham, 2018). These enable to infer DS models, the governing equations underlying experimental data, directly from time series recordings of the system, foregoing any strong a-priori assumptions.

Recurrent neural networks (RNNs) have emerged as particularly powerful deep learning tools in this regard (Fig. 1; (Durstewitz, 2017a; Kramer et al., 2022; Pandarinath et al., 2018; Rajan et al., 2016). RNNs are known to be dynamically universal in the sense that they can approximate any other DS to arbitrary degree (Funahashi & Nakamura, 1993; Hanson & Raginsky, 2020; Kimura & Nakano, 1998; Trischler & D’Eleuterio, 2016). They are also computationally universal in the sense that they can implement the operations of any Turing machine (Chung & Siegelmann, 2021; Hyötyniemi, 1996; Koiran et al., 1994; Sontag & Siegelmann, 1995). RNNs thus possess mathematical properties that make them powerful enough to mimic any neurophysiological or neuro-computational process manifest in the brain. While this is true in theory, progress in RNN training algorithms, especially within the last decade, brought these theoretically pleasing properties also to the ‘workbench’, enabling DS reconstruction from empirical data. Furthermore, from a neuroscience angle, many of the RNN models in use can be thought of as (high-level) abstractions of neural population models. Thereby they offer additional layers of biological interpretability in the neuroscience context, beyond their nature as abstract models of neuronal dynamics (cf. Fig. 10).

**Fig. 1.**
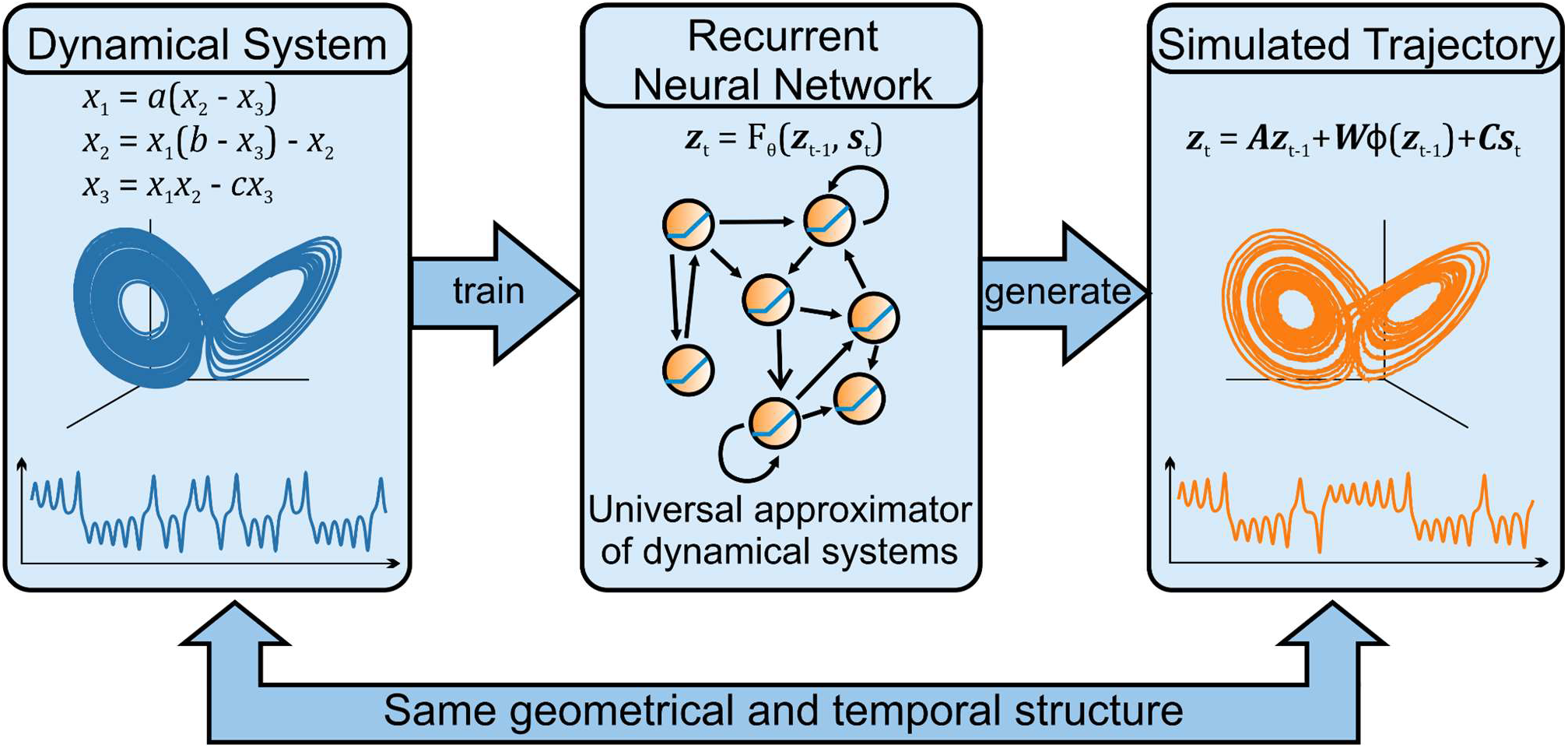
Dynamical Systems (DS) Reconstruction (DSR) via Recurrent Neural Networks (RNNs). In DSR, we train RNNs (center) on time series observations from some (usually unknown) DS (left). After successful training we expect the RNN to generate trajectories and time series with the same geometrical and temporal structure (right) as those produced by the underlying DS the RNN had been trained on.

In this article, we will first provide some important background in DST that helps to understand the virtues of RNNs as theoretical tools in neuroscience (sect. 1). We will then continue with some “historical” notes on *DS reconstruction*, its roots in DST, and the theoretical properties we require in this context (sect. 2.1). This is important to understand when, and under which conditions, we could meaningfully speak of a successful DS reconstruction that permits us to use an RNN as a formal surrogate for the dynamics of empirically observed systems. In our perception, this crucial point is sometimes a bit overlooked in neuroscience: There are certain *formal requirements* that need to be met for something to constitute a proper state space representation of the underlying dynamics. We will then review the field of DS reconstruction by RNNs (sect. 2.2). Here, we will not only cover the progress in computational neuroscience, but also - or in particular - visit recent trends for DS reconstruction in machine learning and nonlinear dynamics that may be less familiar yet to the neuroscience community. Sect. 2.3 gives an overview of RNN *training algorithms*. Training RNNs in general, and for the purpose of DS reconstruction in particular, comes with a number of caveats and issues that need to be taken care of in order to achieve sensible results. Sect. 2.4 will briefly review RNN applications in neuroscience, while sect. 2.5 will discuss how to evaluate successful RNN-based DS reconstructions, before diving into a number of specific examples from a variety of neural measurement modalities, including multiple single unit (MSU) recordings, fMRI and EEG data. Sect. 3 will deal with the all-important issue of RNN analysis and interpretation – after all, we would like to use these systems to gain insight into the dynamical and computational mechanisms underlying our experimental observations. So we wish them to have certain properties that ease analysis and interpretability. Finally, we will conclude with an outlook and discussion of open problems and future directions in this rapidly expanding field (sect. 4).

## 1. Core concepts in DST explained along neuroscience examples

Let us start by introducing some core concepts in DST. The presentation here (as well as some of the material in sect. 2.1) will closely follow a similar introduction provided in Durstewitz et al. (2021).

### 1.1 State space of a dynamical system

To begin with, we feel it is important to emphasize that DST concepts are not merely ‘metaphors’ or analogies for processes in the nervous system, or just one among many competing descriptions of computational mechanisms. DST is a general mathematical language that can be applied to *any* system that evolves across time and space and that can be described by sets of differential (continuous time) or recursive (discrete time) equations. This includes most of the models formulated in computational neuroscience. DST provides a powerful mathematical framework for describing and analyzing dynamical processes in nature that can, in principle, be *empirically observed and measured*. It helps to explain and understand some generic properties of natural systems, like convergence to equilibrium states, multi-stability, chaos, or oscillations and synchrony, under which conditions these phenomena occur, and how they are modulated, created, or destroyed.

DST concepts often have a natural geometrical and topological representation that make them intuitively accessible (Alligood et al., 1996; Strogatz, 2018). At the heart of this is the concept of a *state or phase space*, the space spanned by all the dynamical variables of the system (Fig. 2). Theoretically, this space needs to be *complete* for something to formally constitute a DS (Perko, 2001), in the sense that any vector point in this space contains all information there is about the system’s current state and future evolution. The state space is straightforward to construct if the whole system of governing equations and hence the set of dynamical variables is explicitly given, i.e. if we have a mathematical model of the whole process available. Empirically, if we only have observed a DS through an incomplete and noisy set of measurements, this is much less straightforward. This is exactly the topic of DS reconstruction (DSR) which we will dive into in sect. 2. As one specific example, Fig. 2a provides the state space of a simple 2-variable neuron model, while Fig. 2b shows the state space of a simple neural population model. In the single neuron example (Fig. 2a), a point in state space specifies the current voltage and the state of a recovery variable. In the neural population example (Fig. 2b), each point in state space corresponds to precisely one pair of values for the instantaneous firing rates of an excitatory and an inhibitory neuron population.

**Fig. 2.**
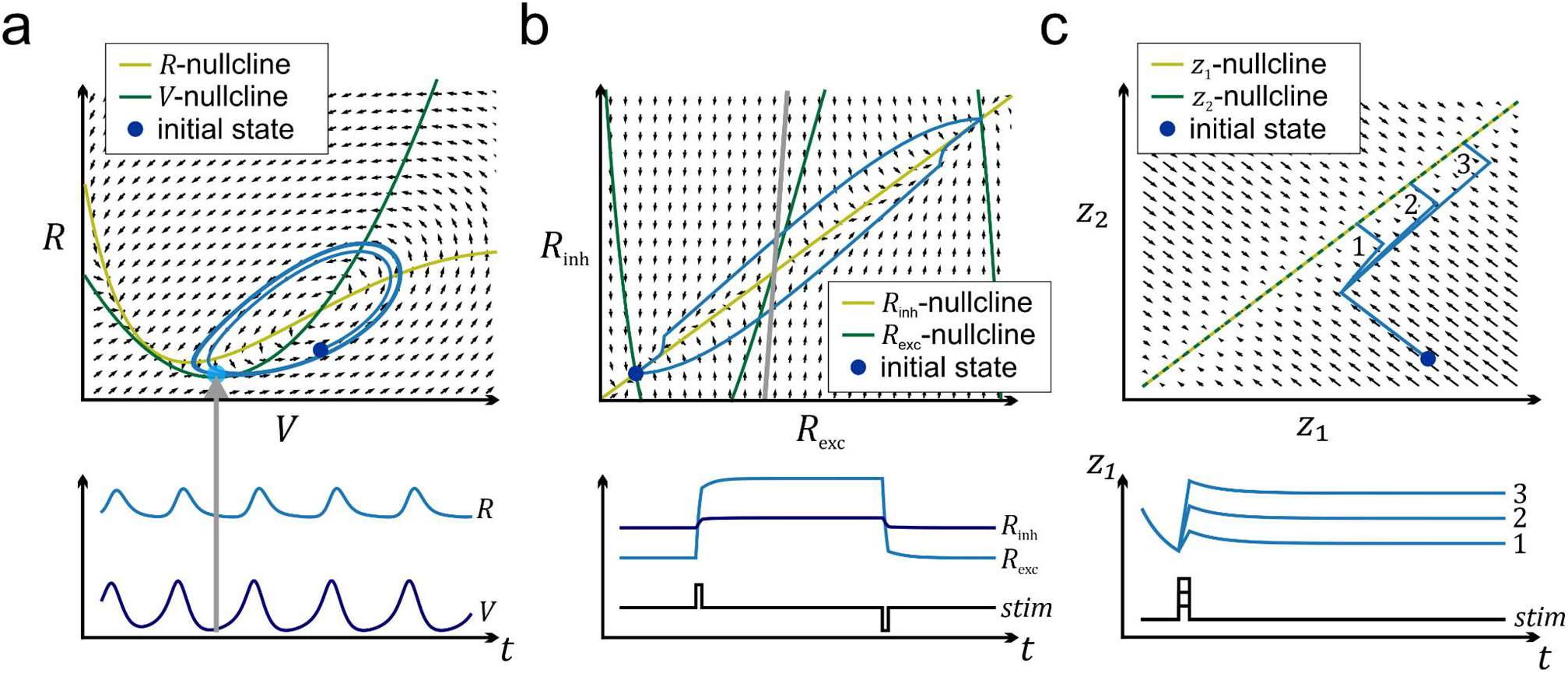
State spaces, vector fields and trajectories. (a) Top: State space with vector field (black) of a 2-dimensional single neuron model exhibiting a limit cycle (regular spiking). Trajectory converging to the limit cycle in blue, with initial condition indicated in dark blue. Nullclines (sets of points where time derivatives for one of the dynamical variables vanish) shown in dark (*V*) and light (*R*) green. Equilibrium (fixed) points are located at intersections of nullclines. Bottom: Temporal evolution of the neuron’s voltage (*V*, dark blue) and recovery (*R*, light blue) variables as the system moves along the limit cycle in state space. Light blue dot indicates position in state space corresponding to the reading of (*V,R*) at the time point given by the gray arrow. (b) Same as (a) for a bistable “working memory” system, given by the firing rates of an excitatory (*R*_exc_) and an inhibitory (*R*_inh_) neuron population. An external stimulus switches the system back and forth between its two stable fixed points, with their basins of attraction (set of initial conditions from which there is convergence to one or the other stable state) demarcated by the gray line. Wilson-Cowan-type model, adapted from Durstewitz (2017b). (c) Illustration of a line attractor configuration: The precise overlap of the system’s two nullclines gives rise to a line of stable fixed points in state space. Starting from the same initial condition, different stimulus strengths drive the system toward different states from where it will converge toward a unique point on the line. It will thereby come to encode a graded memory of the driving stimulus or initial state.

The set of governing equations that mathematically describes (i.e., ‘models’) the system under study gives the precise rules according to which the system evolves in time (and potentially along other dimensions like space). These dynamical rules of the system’s time evolution are geometrically given by its *vector field* (Fig. 2), which prescribes in which direction the system’s state will move at any point in its state space. When started from any such *initial condition*, the system will move through state space in accordance with its vector field, giving rise to a specific *trajectory or orbit* (Fig. 2). Geometrically, each trajectory reflects the joint temporal evolution of the system’s variables, e.g. the neurons’ spiking rates, as depicted in its time graphs. That is, there is a strict 1:1 correspondence of the state of the system’s variables at any point in time and its representation in state space, and exactly one direction of movement associated with each vector point. Formally, a trajectory corresponds to the time solution of the set of differential equations given a specific initial condition (Perko, 2001). The beauty of a state space representation and its vector field is that it yields a compact and complete description and, in low (≤3) dimensions, visualization of the behavior of a DS.

### 1.2 Attractors

A DS’s state space is filled with geometrical objects that determine its behavior in time (and possibly space). The most important class of such objects are *attractors*. Observe in Fig. 2b that the system’s vector field commands convergence of trajectories either to a specific point in the lower left when initiated left from the gray line, or convergence to a specific point in the top right when initiated right from the gray line. These points to which the system’s state converges from some neighborhood, the so-called *basin of attraction* (demarcated by the gray line), are the simplest of all attractor objects, so-called *fixed point attractors* as the limit set to which the trajectories converge consists only of a single point (technically, the term ‘fixed point’ is often reserved for just discrete time systems, while we call them *equilibrium points* in continuous time systems). A simple example of a fixed point attractor in the nervous system is the resting potential of single neuron recorded in isolation: Any (sufficiently small) positive or negative deflection (perturbation) of the membrane potential by a transient current injection will decay back to this stable equilibrium state. Another observation that has been interpreted in terms of fixed point attractors (in firing rates) is that of persistent activity recorded during simple working memory tasks (as in Fig. 2b).

A salient property of the model system depicted in Fig. 2b, reflected in single unit recordings from prefrontal cortex (Funahashi et al., 1989; Fuster, 2015; Fuster, 1973; Miller et al., 1996), is the simultaneous existence of *multiple stable fixed points* (attractors). This phenomenon is called *multi-stability* and is assumed to be of huge functional importance in computational neuroscience. For instance, multiple fixed point attractors may encode different perceptual items to be held active in working memory, with external inputs switching the system between these different states (Fig. 2b; Durstewitz et al., 2000). Or they may correspond to different choice options in a decision making task (Albantakis & Deco, 2009; Wang, 2008). Formally, fixed points are defined by the condition that the vector field (that is all temporal derivatives) becomes exactly zero at these points. *Unstable* fixed points from which the system’s state diverges along one or multiple directions exist as well, as in the center of Fig. 2b (these are *not* attractors, obviously, but are called *repellers, sources*, or *saddle nodes*, depending on their exact properties). Fixed points may also form a continuous line or plane (or any other type of manifold) in state space, a concept denoted *line or plane attractor* (Fig. 2c; Seung, 1996; Seung et al., 2000). It has been hypothesized, for example, to play a role in ‘parametric working memory’ (Machens et al., 2005) where a continuously valued variable like a flutter frequency needs to be maintained (Romo et al., 1999), in maintaining arbitrary eye positions (Seung et al., 2000), in hippocampal head directions cells (Zhang, 1996), in providing context information during decision making (Mante et al., 2013), or in regulating interval timing (Durstewitz, 2003).

### 1.3 Limit cycles and chaos

Attractors do not need to be just single points, but also come in the form of closed orbits, so-called *limit cycle attractors* (Figs. 2a, 3a-1, 3a-3). Stable limit cycles correspond to a nonlinear oscillation in a system: The system’s state continuously cycles along the orbit once on it, and is attracted toward this orbit from some neighborhood (like fixed points, limit cycles can also be unstable or half-stable). Examples of such phenomena in the nervous system are plenty: Regular spiking patterns observed in single neurons constitute stable limit cycles (Izhikevich, 2007), whether rather simple with a single period (Fig. 3a-3) or more complex, multi-period, as in bursting neurons (Fig. 3a-1). Many stereotypical motor or locomotion patterns, like those produced by central pattern generators (Marder et al., 2015; Marder & Bucher, 2001), likely reflect limit cycle attractors as well. Limit cycles have also recently been suggested to underlie the rotational dynamics observed in motor cortex of non-human primates (Lindén et al., 2022; Russo et al., 2018, 2020). Just like with fixed points arranged as line attractors, there are also ‘semi-continuous’ sets of cycles, e.g. as observed in the ‘nearly continuous’ range of stable firing frequencies observed in isolated prefrontal cortical neurons (Fransén et al., 2006; Haj-Dahmane & Andrade, 1999) But stable, attracting activity patterns in the nervous system do not need to be regular at all; they could also be highly *irregular*, as in *chaotic attractors* (Fig. 3a-2). Chaotic attractors correspond to bounded regions in state space that attract nearby trajectories, but with orbits that never precisely close up as in a limit cycle. Evidence for chaotic attractors in neural activity has been obtained both at the single neuron (Durstewitz & Gabriel, 2007) and the network level (Landau & Sompolinsky, 2018; London et al. 2010), and their functional implications for neural computation have been widely discussed (Bertschinger & Natschläger, 2004; Tsuda, 2001, 2015).

**Fig. 3.**
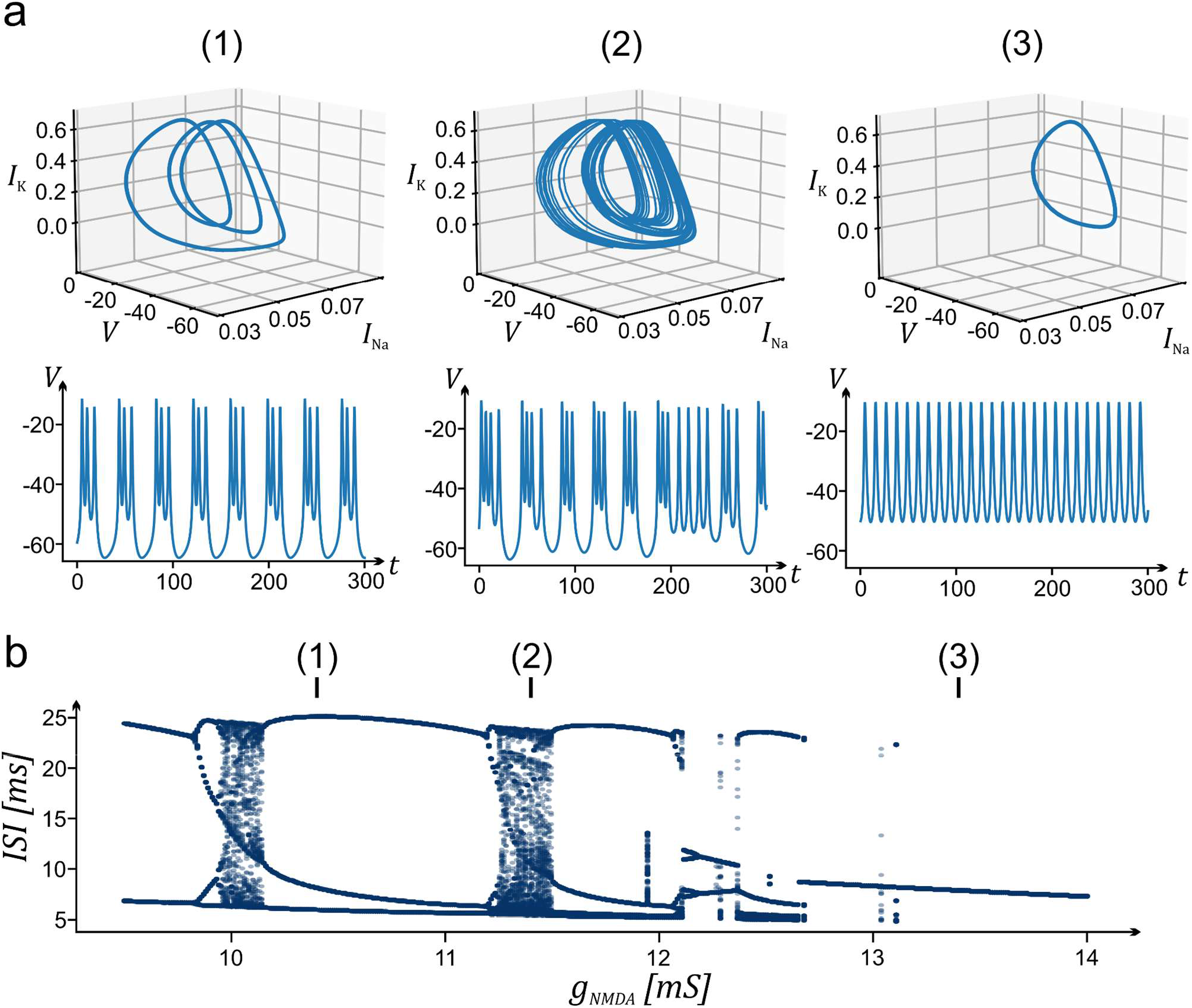
Bifurcations and chaos in a bursting neuron model. (a) State spaces (top) and time graphs of the voltage variable (bottom) of a 3-variable minimal biophysical model of bursting activity induced by NMDA currents (adapted from Durstewitz, 2009). From left to right: Three different levels of NMDA input, resulting in bursting (1), chaotic spiking (2), and regular spiking (3). Similar transitions have been observed in rat prefrontal neurons in-vitro driven by different levels of ambient NMDA (Durstewitz & Gabriel, 2007). (b) Bifurcation graph showing the sets of distinct interspike intervals (ISI) as a function of the NMDA conductance (*g*_NMDA_) control variable. Regimes shown in (a) indicated by the numbers on top.

### 1.4 Bifurcations

Another fundamental concept in DST with huge functional implications is that of a *bifurcation*: Bifurcations denote points (or curves) in parameter space at which *qualitative* changes in the system dynamics occur as its parameters are smoothly varied (Fig. 3). For instance, at a bifurcation point previously stable objects may suddenly lose their stability, novel geometrical objects may come into existence, or others may vanish. When a current injected into a single cell is gradually increased and the cell suddenly starts spiking, then that is an example of a bifurcation (there are a variety of different types of bifurcations which can lead into spiking or bursting activity, each of which endows a neuron with different functional and response properties; see e.g. Durstewitz & Gabriel, 2007; Izhikevich, 2007; Rinzel & Ermentrout, 1998). Sudden transitions in neural population representations during rule learning or shifting tasks (Durstewitz et al., 2010) have also been interpreted as signatures of a bifurcation as a consequence, for instance, of synaptic plasticity associated with acquiring a new behavioral rule. Transitions into and out of psychiatric episodes may also reflect bifurcations (Durstewitz et al., 2021). A crucial point about many types of bifurcations is that they imply an *abrupt change* in system dynamics as a critical point is crossed, a possibility vigorously discussed in the current literature on climate change, for instance (McKay et al., 2022).

There are many other central concepts in DST important to neuroscience, some of which allowing for more flexible structures, as, e.g., required in language, such as *separatrix cycles* (Perko, 2001), also called *heteroclinic channels* (Rabinovich et al., 2008; Rabinovich & Varona, 2011). A comprehensive discussion of DST and its incarnations in neuroscience is far beyond the scope of this review article, but the concepts introduced above should provide sufficient background to appreciate the aims of DS reconstruction discussed next. It may be important to point out that computational neuroscience so far has focused mostly only on the most simple DS phenomena, such as convergence to fixed points, line attractors, or limit cycles. DST provides a much richer world, and so we believe it is fair to say that we have hardly scratched the surface here. For instance, how *flexible computations* as needed, e.g., in problem solving or mental planning, are realized from a DS perspective largely remains a mystery up to now.

## 2. Reconstructing computational dynamics from time series data

### 2.1 Delay coordinate maps and topological equivalence

When we have a mathematical model of a neural process at hand, we can use tools from DST to understand and explain its mechanisms, and to derive functional implications for neural computation (Durstewitz & Seamans, 2002; Wang, 2002; Wilson, 1999). But how can we infer state spaces and their topological and geometrical properties from *experimental data*? There are many statistics, like power laws (Beggs & Plenz, 2003; Jensen, 1998) or change points (Durstewitz et al., 2010; Russo et al., 2021; Toutounji & Durstewitz, 2018), we may interpret as signatures of particular dynamical regimes or events (like bifurcations). To really understand how the brain implements computation, however, a more complete, as well as a more specific, picture of the dynamics underlying experimental measurements is needed. This is the topic of *DS reconstruction*.

A classical tool for reconstructing *trajectories* of an empirically observed DS is the technique of *delay embedding* (Kantz & Schreiber, 2004; Fig. 4a). First, note that we never *directly observe* a DS: We only access quantities like spike times more or less directly related to the underlying DS through some measurement device (like an amplifier), but never the complete set of underlying dynamical variables that give rise to these measurements (like all the ionic currents, or potentially more abstract quantities that exhaustively describe the underlying geometry or topology). Assume you have taken univariate time series measurements *x*_*t*_ from a DS, where these measurements are some function *x*_*t*_ = *h*(***y***(*t*)) of the underlying, unknown DS states ***y*** at time *t* (*h* is also called the observation or measurement function). From these measurements we then form time delay vectors ***x***_t_ by concatenating the measured variables at different time lags, e.g. ***x***_t_ = (*x*_t_, *x*_t-τ_, *x*_t-2τ,_ …, *x*_t-(m-1)τ_), assuming we measured only a scalar quantity, where *m* is the so-called embedding dimension and *τ* is a time-lag (the latter is only practically relevant, not theoretically, and is often chosen such as to sufficiently unfold trajectories without loosing the temporal contiguity between successive data points, Fig. 4a (Kantz & Schreiber, 2004)).This is called a *delay coordinate map*. The delay embedding theorems (Sauer et al., 1991; Takens, 1980) ensure that if *m* is large enough (formally, larger than two times the box-counting dimension of the underlying attractor; Alligood et al., 1996), the reconstructed dynamics will represent the original dynamics in a 1:1 fashion in the sense that all of its topological properties are preserved (Alligood et al., 1996; Sauer et al., 1991; formally, we speak of a diffeomorphism here, a one-to-one map between original and reconstructed state space which is also one-to-one in its derivatives [tangent directions], Fig. 4b). Delay embedding reconstructions were quite popular in physics in the 90es-2000s (Kantz & Schreiber, 2004). They have also been employed in neuroscience for reconstructing neural semi-stable states during multiple-item working memory tasks from MSU recordings (Balaguer-Ballester et al., 2011). Noise, of course, may diminish our ability to reconstruct the underlying system fully, but that is a longer discussion in its own right.

**Fig. 4.**
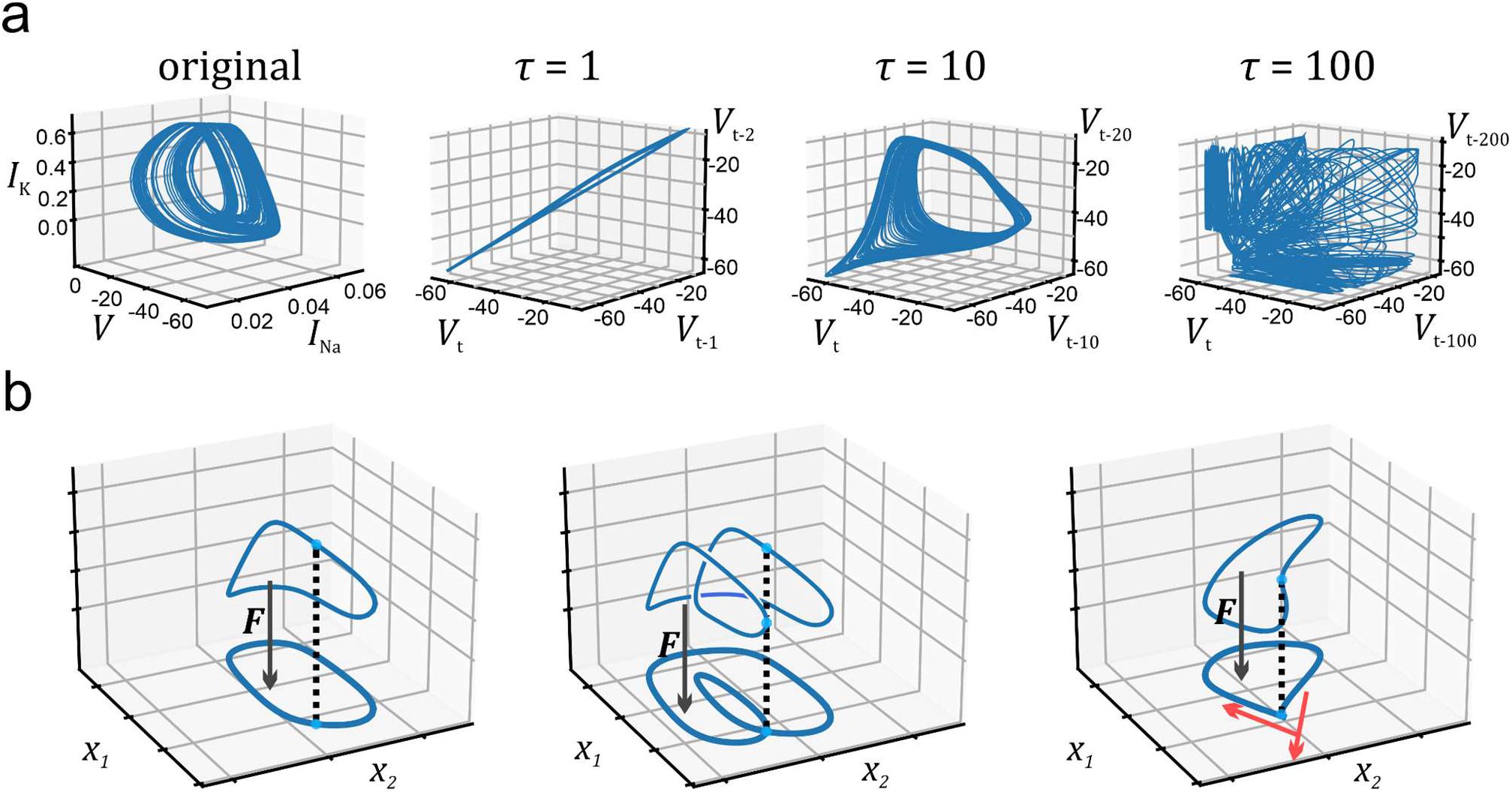
Delay embedding and topological equivalence. (a) Delay coordinate embedding of trajectories of the bursting neuron model on the left. Shown are embeddings (*V*_*t*_, *V*_*t*-τ_, *V*_t-2τ_) for three different time lags *τ* of the neuron’s voltage variable. Choosing *τ* too small will lead points to cluster along a thin line, while choosing *τ* too large will result in very erratic behavior as successive time points become essentially unrelated (at least in the presence of noise). (b) Examples of topologically equivalent mapping *F* of a limit cycle trajectory to a 2d space (left), violation of 1:1 property of map *F* (center), and violation of 1:1 property of tangent map *F*′ (first derivatives, collected in the so-called Jacobian matrix). Adapted from Sauer et al. (1991), using the neuron model from Fig. 3.

There are important mathematical insights behind delay reconstructions: In any empirical situation, we can never be sure that all relevant degrees of freedom of the DS under study have been properly probed. The delay embedding theorems give a guideline of how we need to augment the space empirically accessed to be sure that dynamical objects in the reconstructed space topologically correspond to those in the original DS in a unique fashion. Pleasingly, delay embedding theorems also exist for spike train data, which are formally point processes (Sauer, 1994, 1995). Crucially, simple dimensionality reduction tools like PCA as commonly employed to represent ‘neural state spaces’ *do not share these properties* and theoretical guarantees. In fact, they may even destroy important dynamical features, since they form linear combinations of observations that may not distinguish between different states in the system’s true state space (i.e., do not come with the required 1:1 properties). At the very least, unlike assured by the delay embedding theorems (Sauer et al., 1991), they come with no theoretical guarantees that the system dynamics is properly conserved and represented in the reduced spaces, and this is not what they were designed for. The same holds true for any of the more recent nonlinear dimensionality reduction tools like Locally Linear Embedding (Roweis & Saul, 2000), Isomap (Tenenbaum et al., 2000), Laplacian eigenmaps (Belkin & Niyogi, 2003), t-SNE (Van der Maaten & Hinton, 2008), or autoencoders (Hinton & Salakhutdinov, 2006), for instance.

Delay-coordinate maps only reconstruct the particular attractor object probed by the experimental measurements. They provide an embedding for the specific experimental trajectories observed, but do not yield a full reconstruction of the underlying DS, i.e., the governing equations. For testing ideas about neural computation, we would like to have more information about other attracting or unstable objects, their specific geometrical and stability properties, their basins of attraction, stable and unstable manifolds, vector fields throughout state space, and so on. Ultimately, we would like to have a *mathematical model* of the system under study at our hands that we can scrutinize and analyze in depth. In fact, if we are interested in *computational* properties of neural activity, then – quite tautologically – we need a computational model.

One idea to approach this goal is to formulate a mathematical model based on first principles, e.g. a biophysical model of neural systems, whose parameters we then try to infer from the experimental data (Abarbanel, 2013; Hass et al., 2016; Lueckmann et al., 2017; Raissi et al., 2018). A potential disadvantage here is that we need to specify the full form of the underlying model (the complete system of equations) in advance. In doing so, we need to make potentially rather strong assumptions about the underlying biophysics. Such ‘structural priors’ may be helpful if the data are rather sparse, but they entail a risk of severe model misspecification (‘bias’). Especially in areas where the systems are very complex and knowledge is still rather sparse (as in neuroscience), this may easily result in a model with dynamics quite different from that of the experimentally probed system (although there are diagnostic checks one can do, see Abarbanel, 2013). For instance, if we ‘forget’ to include a dendritic voltage-gated Ca^2+^ channel in our biophysical model, our inference scheme would be forced to attribute its effects to other biophysical mechanisms. This in turn may mess up the model’s dynamical properties in profound ways. Perhaps more importantly, complex models based on first principles are also often highly challenging to analyze in themselves.

### 2.2 Machine learning models for dynamical systems reconstructions

A quite different approach to DS reconstruction is to set up a mathematical model which is a *universal function approximator*. By this we mean a set of equations which is powerful and expressive enough to approximate any other function to arbitrary precision (provided some mathematical conditions are met). Polynomial basis expansions are known to have this property (by which we mean there are mathematical theorems which assure this: e.g the Stone-Weierstrass approximation theorem; Llavona, 1986). Neural networks with at least one nonlinear hidden layer also fall into this class (Cybenko, 1989; Hornik et al., 1989; Lu et al., 2017). Hence, in theory, we could use a basis expansion (Brunton et al., 2016; Storace & De Feo, 2005) or a neural network (Chen & Chen, 1995; Hanson & Raginsky, 2020; Kimura & Nakano, 1998; Trischler & D’Eleuterio, 2016) to closely approximate the vector field of any given DS, thereby, ideally, reconstructing its topological and geometrical properties in full (Fig. 5). At the same time, it may be important to emphasize that there are many popular ML models which do *not* enjoy these mathematical properties. *Linear* dynamical systems, for instance, like linear state space models (e.g., Friston et al., 2003; Macke et al., 2015; Paninski et al., 2010; Yu et al., 2009), are *inherently* (mathematically) incapable of producing most of the phenomena discussed in sect. 1, including limit cycles, multi-stability, separatrix cycles, or chaos (Perko, 2001; Strogatz, 2018).

**Fig. 5.**
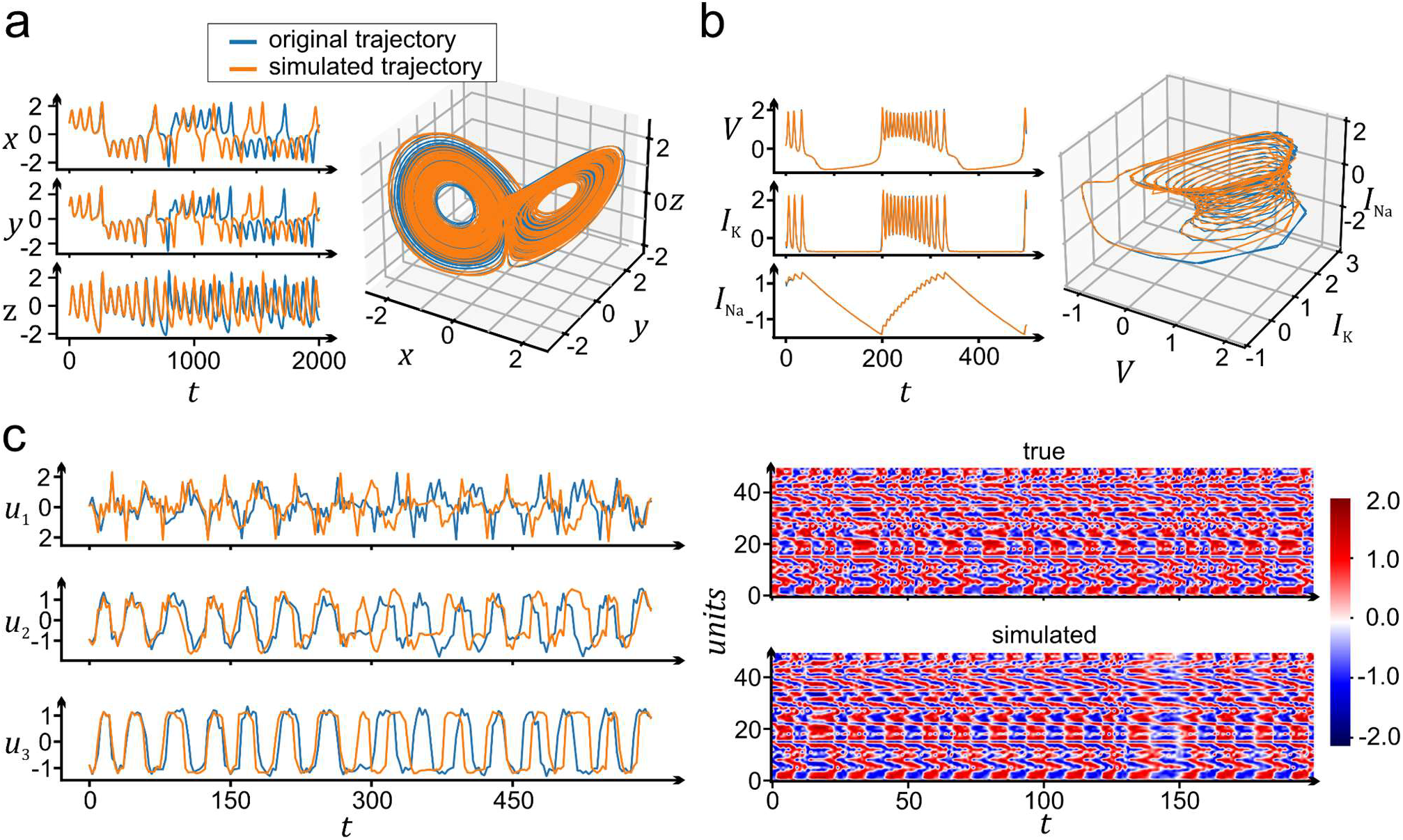
Reconstruction of benchmark DS by RNNs. (a) True (blue) and RNN-generated (orange) trajectory as time graphs (left) and in state space (right) for the Lorenz-63 weather system. For producing the orange trajectory, a trained RNN was released from an initial condition inferred from the data and then freely forward-iterated in time (i.e., without any further reference to the data previously used for training). Note that due the chaotic nature of the system, true and RNN-simulated trajectories tightly agree only for some hundred time steps, but then start to diverge, yet – and importantly – retain the same temporal and geometrical structure. (b) Same as (a) for a biophysical neuron model in a bursting (non-chaotic) regime. (c) Same for a high-dimensional system (neural population model by Landau & Sompolinsky, 2018). Left shows some selected units, while the right panel shows the whole spatio-temporal pattern. All graphs based on Brenner et al. (2022).

Here is another important point: The mathematical form of the equations used for approximation can be *completely different* from those that most “naturally” describe the true DS. For instance, in Fig. 5b, a piecewise-linear recurrent neural network (PLRNN), containing only piecewise-linear terms, has been trained to approximate the dynamics of a biophysical spiking neuron model (which consists of equations containing exponential and multinomial terms). Thus, as highlighted by this example, *we do not need a detailed biological model specified by biophysical equations to perfectly mimic the behavior of a biological neuron*. This is a truly remarkable fact – in theory, no prior knowledge about the DS in question is needed. Whatever it is, we know it can, in principle, be faithfully reproduced, with all its detail, by a generic set of RNN or basis expansion equations just trained on observations from the true system. In practice, however, good prior knowledge about the governing set of equations can help to lower the data requirements, and, hence, some ML approaches attempt to combine structural priors with pure data-driven ‘black box’ models (Haußmann et al., 2021; Singh et al., 2019; Yin et al., 2021).

Within the last decade or so, a large variety of ML approaches for DS reconstruction based on basis expansions (Brunton et al., 2016; Champion et al., 2019), Fourier neural operators (Li et al., 2021), Koopman operators (Lusch et al., 2018), Transformers (Shalova & Oseledets, 2020), Graph Neural Networks (Iakovlev et al., 2021), continuous-time (Haußmann et al., 2021) or discrete-time RNNs (Brenner et al., 2022; Koppe et al., 2019; Mikhaeil et al., 2022; Seleznev et al., 2019; Vlachas et al., 2018, 2020) has been introduced in machine learning, nonlinear dynamics, and physics; too numerous to be all reviewed in this short introduction. The focus here will be on RNNs, which have become highly popular recently in neuroscience (Barak, 2017; Hernandez et al., 2020; Kim et al., 2021; Koppe et al., 2019; Kramer et al., 2022; Mante et al., 2013; Nayebi et al., 2021; Pandarinath et al., 2018; Rajan et al., 2016; Sussillo & Abbott, 2009; Wang et al., 2018; Yang et al., 2019). An RNN is essentially a high-level abstraction of a neural system that combines a linear transformation of a neuron’s ‘synaptic’ inputs with a nonlinear activation function which yields its ‘firing rate’ output (Fig. 1). RNNs differ from feedforward networks, like convolutional neural networks, as most commonly used in machine learning, by the presence of recurrent connections which allow for the ‘reverberation’ of activity within the network. Formally, this makes RNNs DS by themselves. It enables them to both process incoming time series as well as produce diverse temporal patterns, as for instance in natural language processing and machine translation (Otter et al., 2021; Yu et al., 2019). A great variety of different RNN architectures has been proposed since the 90es, some formulated in discrete time (i.e., as time-recursive equations; e.g. Long-Short-Term Memory [LSTM; (Hochreiter & Schmidhuber, 1997)] or Gated Recurrent Units [GRUs (Cho et al., 2014; Chung et al., 2014)], some in continuous time (i.e., as systems of ordinary or partial differential equations; e.g. Chen et al., 2018; Rusch et al., 2022; Rusch & Mishra, 2021a; Salvi et al., 2022; Song et al., 2016). RNN-driven *DS reconstruction* methods, in particular, have been based, for instance, on Long-Short-Term-Memory (LSTM) networks (Vlachas et al., 2018), Reservoir Computing [RC; (Jüngling et al., 2019; Pathak et al., 2018)], ODE/PDE-based RNNs (Haußmann et al., 2021; Salvi et al., 2022), or piecewise-linear RNNs [PLRNNs; (Brenner et al., 2022; Durstewitz, 2017a; Koppe et al., 2019; Schmidt et al., 2021].

Let us take a closer look at a PLRNN, for example, defined as follows (Koppe et al., 2019; Schmidt et al., 2021):

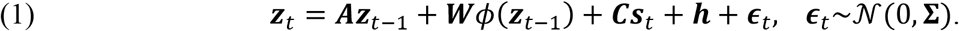

Here, ***z***_t_ is a vector-valued variable (the RNN’s latent states) that evolves in time according to the dynamical rule given by eq. (1), and ***∈***_t_ is a Gaussian noise term. The diagonal entries in ***A*** one may interpret as the units’ passive time constants, the entries in matrix ***W*** as synaptic coupling terms, the so-called rectified linear unit (*ReLU*) operation *ϕ*(***z***) = max(***z***, 0) as a nonlinear voltage-to-spike-rate transfer function, ***h*** as a base rate (or constant/ mean input), and ***s***_t_ as external inputs to the system weighted by synaptic weights ***C***. A PLRNN may thus be seen as a simple discrete-time neural population model (the same is true for some other RNN formulations), although not important for its application in DS reconstruction.

We close this section by pointing out that there are also RNN-based reconstruction algorithms that allow for training on *multiple data modalities* simultaneously (Kramer et al., 2022). This works by coupling the same underlying RNN latent model to several so-called modality-specific observation (or decoder) models which capture the unique statistical properties of each data modality observed (Fig. 6). For instance, in neuroscience we often have simultaneous recordings of neural activity and behavior, where the latter often consists of categorical random variables (e.g., left vs. right lever presses in a typical experiment). Integrating different data modalities into the same latent RNN for DS reconstruction enables one to directly examine their relations in the RNN’s latent space. For example, simultaneous training on neuronal and behavioral data could establish direct links between neural trajectories, DS objects, and behavioral choice processes (Kramer et al., 2022).

**Fig. 6.**
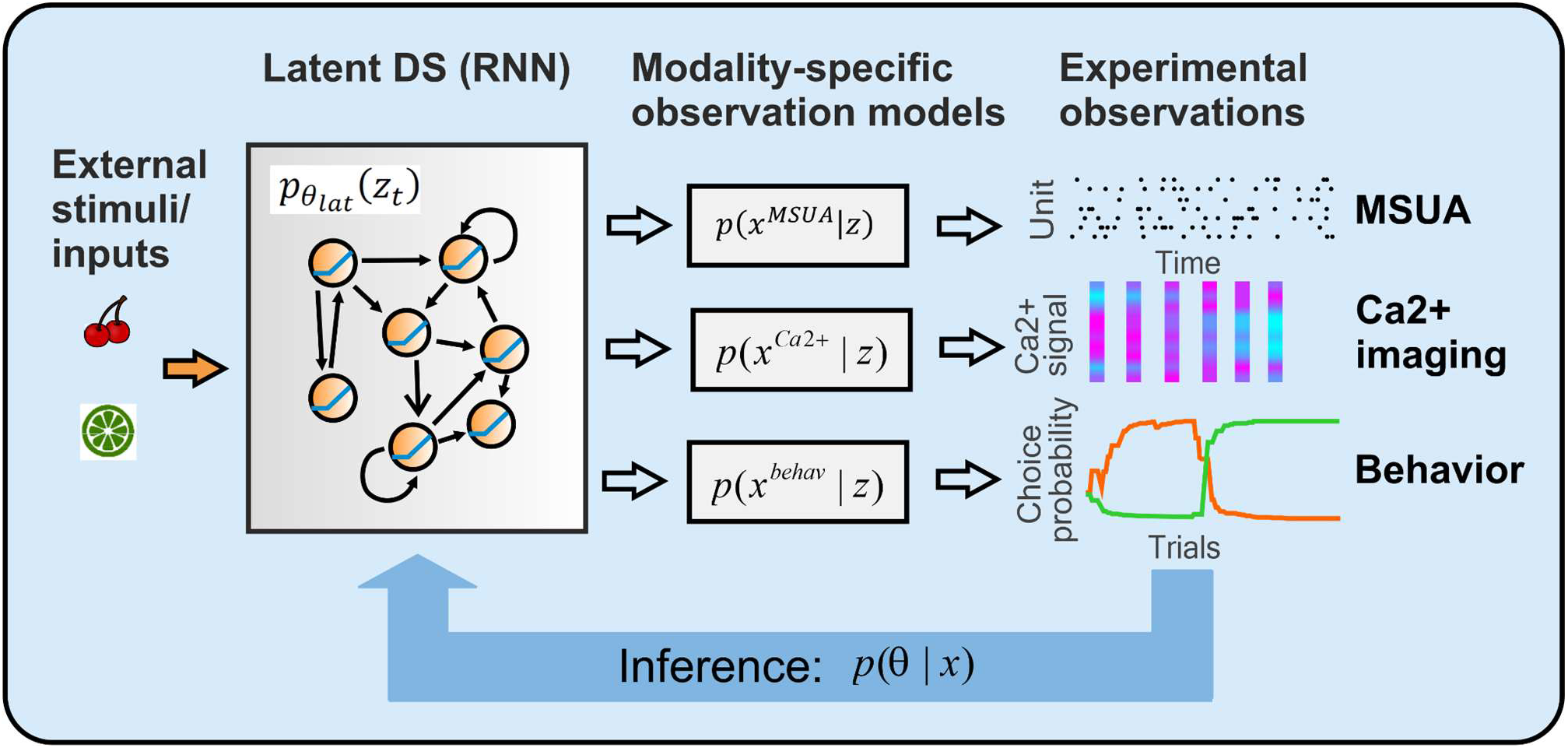
DS reconstruction from multi-modal time series data. Different types of simultaneously recorded time series data with diverse statistical properties, e.g. from categorical (like behavioral), count (like spike counts) or continuous-Gaussian (like neuroimaging) processes, may be coupled to the same latent RNN through a set of modality-specific observation models. Such a framework gives rise to a joint multi-modal likelihood allowing for inference of model parameters simultaneously from different types of datasets. Modified from (Kramer et al., 2022).

### 2.3 Training RNNs on dynamical systems

Training algorithms for RNNs may roughly be distinguished by whether they are probabilistic or deterministic, and by whether they are of first or higher order. *Probabilistic* RNN training algorithms like Expectation-Maximization (Durstewitz, 2017a; Koppe et al., 2019; Voss et al., 2004) or variational inference (Hernandez et al., 2020; Kramer et al., 2022) treat the RNN’s latent states explicitly as random variables and employ maximum-likelihood or Bayesian inference. *Deterministic* algorithms, on the other hand, do not make such explicit assumptions. Usually they attempt to simply minimize the mean-squared-error between observed and predicted outputs. They are far more common and include the famous back-propagation–through-time (BPTT; Rumelhart et al., 1986) and real-time-recurrent-learning (RTRL; Pearlmutter, 1990; Williams & Zipser, 1989) algorithms, which are *first order* gradient descent methods. *Second-order* algorithms instead implicitly or explicitly also consider second or higher order derivatives of the loss function, like recursive-least-squares (Mandic & Chambers, 2001; Sussillo & Abbott, 2009), Hessian-based (e.g., Newton-Raphson), conjugate direction or quasi-Newton methods (Luenberger & Ye, 2016), or “Hessian-free” (Martens & Sutskever, 2011) algorithms. They are in general numerically more precise, but computationally far more costly. All of these have been used in the context of DS reconstruction, but at the moment it is not quite clear whether more complex probabilistic algorithms actually outperform simpler gradient descent methods (e.g. Brenner et al., 2022).

From the discussion in the previous two sections it should already be clear that certain conditions need to be met for a system to yield a proper DS reconstruction. Here we will bring up a few additional points to be considered in practice. In DS reconstruction we are interested in reconstructing the original system’s complete vector field (or at least its topology), rather than just finite-length trajectories (Kimura & Nakano, 1998; Trischler & D’Eleuterio, 2016). In reality, however, we do not have access to the complete vector field of the DS of interest, but usually only to (partial) time series measurements from it. Hence, we either would need to first obtain an estimate of the vector field by numerical differentiation (Brunton et al., 2016; Chen et al., 2017; De Feo & Storace, 2005). Or we could try to infer the model from the time series directly (Durstewitz, 2017a; Hernandez et al., 2020; Koppe et al., 2019; Pathak et al., 2018; Schmidt et al., 2021). Both is possible, but numerical derivatives are usually more noisy and require additional steps for noise reduction (Baydin et al., 2018; Raissi et al., 2018). This issue may be aggravated in very high-dimensional spaces like those we typically encounter in neuroscience, as the observed trajectories would usually sample only a tiny portion of the system’s vector field. Another more general issue to consider is that even if we had available measurements from hundreds to thousands of neurons, there is still no guarantee that all relevant degrees of freedom needed to properly describe the underlying dynamics have been assessed. Thus, delay embedding (see sect. 2.1) may still be necessary, potentially followed by some *dynamics-preserving* form of nonlinear dimensionality reduction to extract the manifold optimally representing the system dynamics (Balaguer-Ballester et al., 2011; Champion et al., 2019; De Feo & Storace, 2005; Lusch et al., 2018). Alternatively, *latent variable models* (i.e., including variables not directly tied to the observations) may be used that harbor sufficiently many degrees of freedom to resolve partially unobserved DS (Hernandez et al., 2020; Koppe et al., 2019; Kramer et al., 2022; Voss et al., 2004).

Training RNNs on observations from chaotic DS, or DS that evolve on many different time scales, poses particular problems. One common issue here is the so-called ‘exploding-&-vanishing gradient problem’ (Bengio et al., 1994; Hochreiter, 1991): Loss gradients in typical gradient-descent-based RNN training algorithms such as BPTT (Rumelhart et al., 1986) tend to quickly decay away or blow up along longer training sequences. This makes it very hard to train vanilla RNNs on time series that involve temporally widely separated events or processes that evolve very slowly, on top of much faster dynamics (see Schmidt et al., 2021). Classical solutions like LSTMs (Hochreiter & Schmidhuber, 1997) or GRUs (Chung et al., 2014) often involve purpose-tailored architectures. More recent approaches try to address the problem through specific architectural (Rusch & Mishra, 2021a, 2021b) or parameter (Arjovsky et al., 2016; Chang et al., 2019; Helfrich et al., 2018) constraints. It is important to note, however, that the resulting systems are often *not suitable for DS reconstruction* as these same constraints rule out (mathematically) many important dynamical phenomena like multistability or chaos (Mikhaeil et al., 2022). In fact, an important insight is that for chaotic systems exploding gradients *cannot be avoided even in principle* (Mikhaeil et al., 2022), as they are a consequence of the exponentially diverging trajectories in chaotic systems. This is a serious issue since chaotic dynamics is rather the rule than the exception in most complex biological or physical systems (see Mikhaeil et al., 2022; Degn et al., 2013). Control-theoretic methods like “teacher forcing” (Pearlmutter, 1990; Williams & Zipser, 1989) that pull diverging trajectories back on track while training, at intervals determined by the system’s Lyapunov spectrum, may offer a remedy (Mikhaeil et al., 2022). Specific optimization criteria that encourage the mapping of both fast and very slow time scales while retaining the structural simplicity of vanilla RNNs have furthermore been introduced to alleviate the exploding-&-vanishing gradient problem specifically in the context of DS reconstruction (Schmidt et al., 2021).

### 2.4 RNNs in neuroscience

So far, RNN models in neuroscience have mostly been used either for posterior inference of neural trajectories in reduced latent spaces (Pandarinath et al., 2018; Kim et al. 2021; Zhao & Park, 2020), i.e. for inferring the most likely latent trajectory given the data. Or they were employed as tools for generating hypotheses about the neural dynamics underlying performance in simple behavioral tasks (Mante et al., 2013; Song et al., 2016; Wang et al., 2018; Yang et al., 2019). For the latter purpose, RNNs are simply trained on the same cognitive or perceptual tasks as used in the animal experiments, and their dynamics are interpreted post-hoc in relation to the neural recordings. This has led to many interesting insights into the potential dynamics underlying cognitive operations, but it is not the same as DS reconstruction, of course. In DS reconstruction we are aiming for more (Figs. 4-5): We would like to obtain an approximation to the *governing equations* of the DS underlying the observed data that shares all topological, and ideally also geometrical, properties with the original system (i.e., produces a 1:1 mapping of the true system’s trajectories and tangent spaces across the *whole* state space). If this were accomplished, the resulting RNN model could be used as a surrogate for the experimentally observed system: From the DS point of view, it would behave equivalently to the underlying system it had been trained on.

Various authors have indeed trained RNNs directly on neurophysiological data (Hernandez et al., 2020; Kim et al., 2021; Koppe et al., 2019; Kramer et al., 2022; Rajan et al., 2016). Again, this may serve multiple purposes other than DS reconstruction in the strict sense. For instance, we may simply take RNNs as model-based nonlinear tools to quantify interactions and information flow among brain areas (see sect. 3), or we may utilize them to analyze the relation between neural and behavioral variables (Kramer et al., 2022). While these are all fair and useful applications, if we are aiming for DS reconstruction, we need to demonstrate more directly that the employed models and training procedures are capable of reconstructing invariant (geometrical and temporal) properties of an unknown data-generating DS (Hernandez et al., 2020; Koppe et al., 2019; Kramer et al., 2022), as will be discussed next.

### 2.5 Evaluating the quality of DS reconstructions

How do we evaluate whether the DS reconstruction worked as supposed? In machine learning, RNNs are mostly used for ahead-predictions on the system under study, e.g. for predicting electricity consumption (Rahman et al., 2018) or forecasting object trajectories (Kim et al., 2017). Mean-squared prediction errors (MSPE) are therefore often used to evaluate the performance of an RNN training algorithm. However, MSPEs are not a sufficient (or even suitable) metric for the evaluation of DS reconstruction algorithms. If the underlying system is chaotic, due to the exponential divergence of nearby trajectories, minuscule amounts of noise or differences in initial conditions will quickly lead to large MSPEs *even if the observations were drawn from the exact same underlying system with the same parameters* (Fig. 7; Koppe et al., 2019). Vice versa, a comparatively low MSPE may falsely indicate a good agreement between a true and a reconstructed system although the two systems may profoundly differ in their underlying dynamics. This could be the case, for instance, when the reconstructed system exhibits a limit cycle with roughly the same dominant frequencies as those of an underlying chaotic system (Fig. 7b, left). In DS reconstruction, it is therefore important to check for geometrical and other time-invariant properties of the systems under study. For instance, measures based on the Kullback-Leibler divergence (Brenner et al., 2022; Koppe et al., 2019) or Wasserstein distance (Patel & Ott, 2022) have been used to assess the geometrical overlap in data point distributions in the large time limit across state spaces generated by the true and the reconstructed system. The maximal Lyapunov exponent or the so-called correlation dimension, an empirical estimate of the fractal dimensionality of an attractor, are other examples of such invariant measures (Kantz & Schreiber, 2004; see also Gilpin, 2020). Agreement in the temporal structure of true and reconstructed trajectories may be assessed by auto-correlation functions or overlap in power spectra (Brenner et al., 2022; Koppe et al., 2019; Vlachas et al., 2018; Wood, 2010), quantified for instance through the Hellinger distance (Mikhaeil et al., 2022), rather than by overlaying time series directly.

**Fig. 7.**
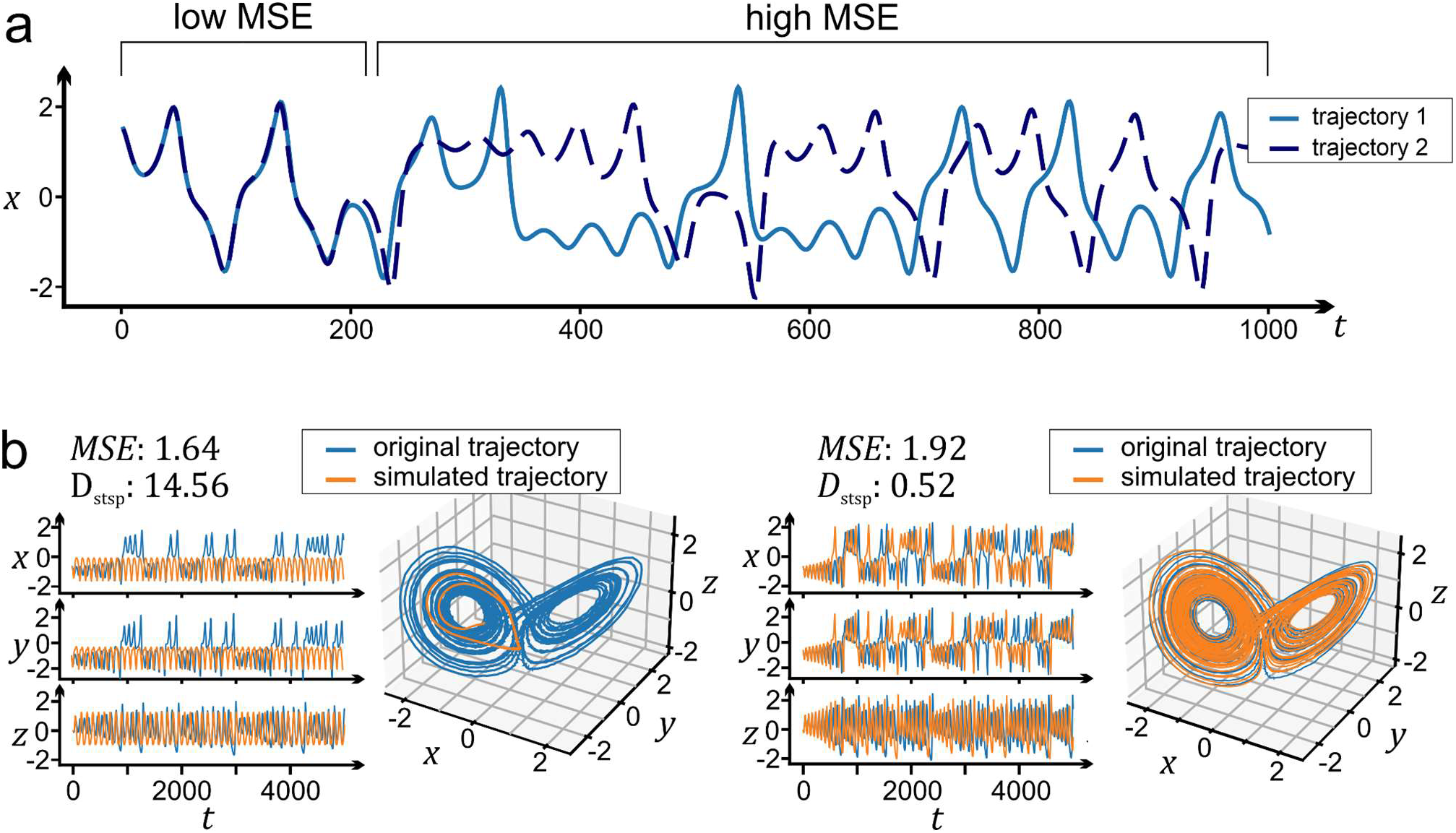
How to measure DS reconstruction quality? (a) The standard ML metric, mean-squared-error (MSE), is not useful beyond a couple of time steps in chaotic systems, as two trajectories drawn from the *very same* system, yet with slightly differing initial conditions, will eventually diverge and produce a large MSE. (b) Due to this fact, a reconstructed system may have a relatively low MSE yet produce very different dynamics as measured through the geometrical overlap (*D*_stsp_) between true and model-generated trajectories (left). This can happen, for instance, if the reconstructed system exhibits a regular oscillation with the same dominant frequency as the chaotic dynamics of the true system. Vice versa, a very well reconstructed system with the same dynamics and geometry as the ground truth system may exhibit a comparatively high MSE (right). Adapted from Koppe et al. (2019).

Ultimately, if the DS reconstruction were successful, we would expect that by *simulating* the RNN we would obtain trajectories with the same temporal and geometrical properties as those produced by the original system. This is illustrated in Fig. 5 for a PLRNN trained on a variety of low- and high-dimensional ground truth systems (Brenner et al., 2022; Schmidt et al., 2021). Note that this is very different from just “fitting” a given data set: The model-generated trajectories in Fig. 5 are obtained by *freely running* the PLRNN from some initial condition, i.e. by evolving its dynamical equations *independently* from the data previously used for training. Thus, after training, the model’s governing equations produce the same dynamics as that of the corresponding ground truth system, and thereby become a surrogate for the true system.

Fig. 8 illustrates such reconstructions for the PLRNN on a variety of real experimental data: In Fig. 8a, fMRI BOLD signals simulated by a PLRNN overlaid with the true BOLD activity are shown, with overlaid power spectra of true and PLRNN-simulated fMRI activity in the center of Fig. 8a (Koppe et al., 2019; Kramer et al., 2022). Fig. 8b shows true and reconstructed spatio-temporal EEG patterns (Brenner et al., 2022). For Fig. 8c, a PLRNN was trained on multiple spike train data recorded from rodent prelimbic cortex during a rule shift task. After training, the PLRNN generates spike rate curves capturing the whole variety of measured neuron’s rate profiles on single trials.

**Fig. 8.**
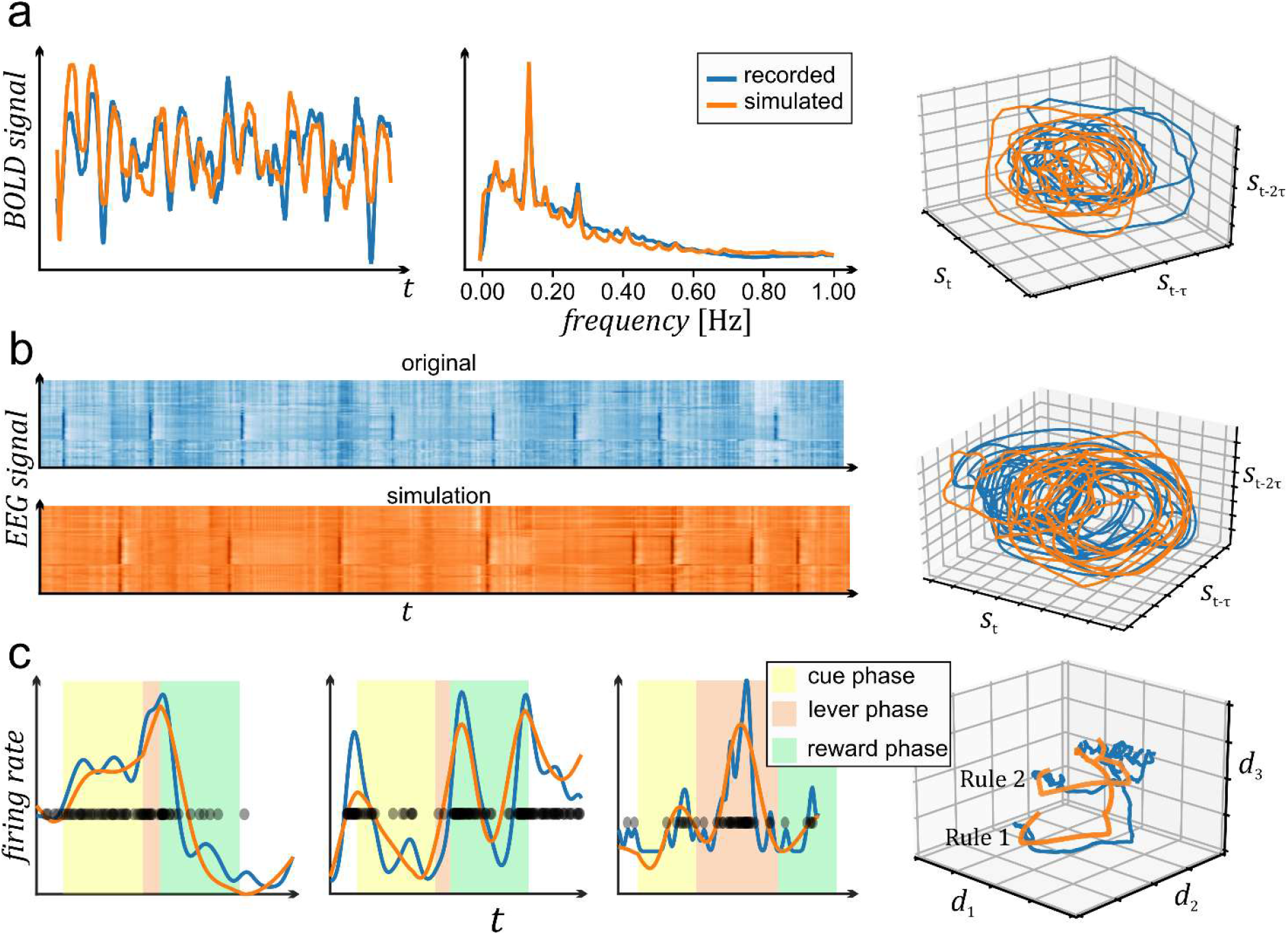
Example reconstructions of real physiological data. (a) Overlaid true (blue) and RNN-generated (orange) BOLD time series (left), power spectra (center), and delay embedding spaces (right). As for the benchmark DS in Fig. 5, RNN-generated trajectories were produced by freely simulating the once trained system. Adapted from Kramer et al. (2022). (b) Same for human EEG data, with overlaid time series from 64 channels on left and delay embedding spaces on right. Data from Schalk et al. (2004), provided at https://physionet.org/content/eegmmidb/1.0.0/. Simulations were generated by Lukas Judith using code from Brenner et al. (2022). (c) Same for single unit data recorded from the rat prefrontal cortex during a rule-shift task (data kindly provided by Dr. Florian Bähner, CIMH Mannheim). Spike trains were first convolved with Gaussian kernels to yield an estimate of instantaneous firing rate, and real and RNN-simulated rates are shown on top of each other for various single unit examples, together with task events, during *single* trials (no averaging). There is a close agreement between model-generated and true unit activity for a wide variety of single unit behaviors. Right panel shows a *projection* of true and RNN-generated MSU trajectories into a lower-dimensional space obtained by Isomap.

Using RNNs for DS reconstruction, as defined here, on multiple single unit recordings, a more complex picture of neural dynamics may emerge than suggested by the effectively low-dimensional, simple ‘fixed point attractor’ models and dimensionality reduction techniques most popular in neuroscience: For instance, as illustrated in Fig. 9, during the life time of a single trial only transient dynamics may be observed (and embody the essence of the computations; cf. Durstewitz & Deco, 2008), while ultimately neural activity may converge to complex chaotic attractors (see also Pereira-Obilinovic et al., 2021).

**Fig. 9.**
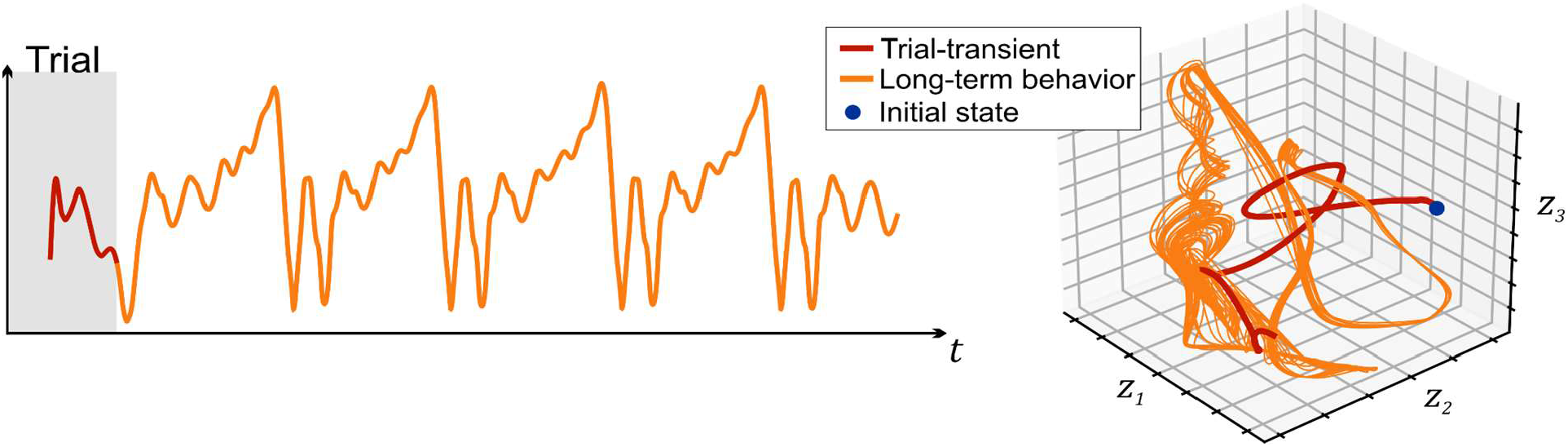
Transient dynamics may rule neural activity on single trials, not the attractor states the activity may ultimately converge to. In this example, an RNN trained on multiple single unit recordings from the rodent prefrontal cortex (see Fig. 8) was simulated beyond the length of a typical trial, as indicated by the gray box and change in coloring in the time graph (left) and a 3d subspace of the RNN’s state space (right). During the duration of the actual trial the trajectory never reaches any attractor state, while ultimately (beyond the trial) it converges into a more complex chaotic attractor.

## 3. Analysis and interpretation of trained RNNs

Suppose we have confirmed an RNN is capable of generating new trajectories with the same temporal and geometrical properties as those exhibited by the original system it has been trained on. How do we proceed from there? Generally, any data-inferred RNN model offers two related levels of interpretability: First, we can inspect the model’s parameters to infer physiological or anatomical properties of the underlying system, e.g. connectivity between neurons or brain areas (Fig. 10). This we may do, in principle, without proper DS reconstruction (although it would certainly help to lend credit to the results). Second, we can use DST tools to analyze in depth the dynamical properties of the model in order to reveal how the underlying neuronal substrate performs the computations necessary to accomplish the behavioral task at hand (Fig. 11). Both these incentives will be briefly discussed below. Of course, it is the latter purpose which we are mostly interested in here and where data-inferred RNNs become particularly powerful.

**Fig. 10.**
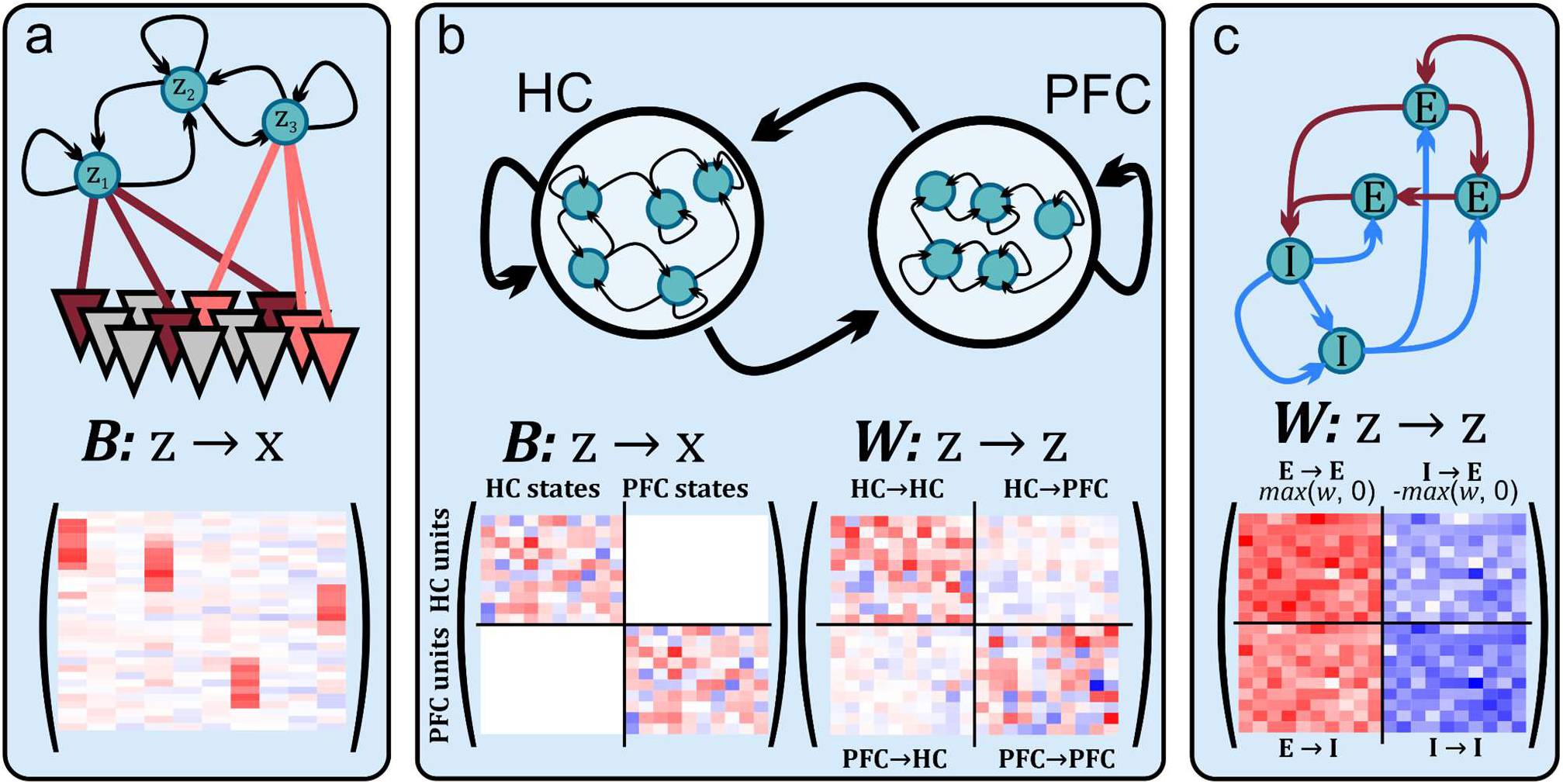
Interpretation of trained RNNs: Relation to biological substrate. Structure in the observation matrix ***B*** (cf. eq. 2) and the connectivity matrix ***W*** (cf. eq. 1) among latent states enables to link dynamics to biological underpinnings. (a) Different recorded units loading on the same latent state may be interpreted as part of the same assembly (see also Fig. 11b below). (b) By constraining the observation matrix ***B*** such that subsets of RNN latent states can couple to recorded neurons only from one specific brain area, the connectivity matrix ***W*** acquires biological meaning in terms of *within-vs. between area connections*. This can be used to examine inter-areal information transfer in a time-resolved manner. (c) Likewise, ***B*** may be structured such that different latent states couple only to specific neuron types (e.g., pyramidal vs. interneurons), and their weights can easily be constrained through the *max* operator to be only excitatory (positive) or inhibitory (negative). This way, the role of different neuron types in the dynamics could be dissected.

**Fig. 11.**
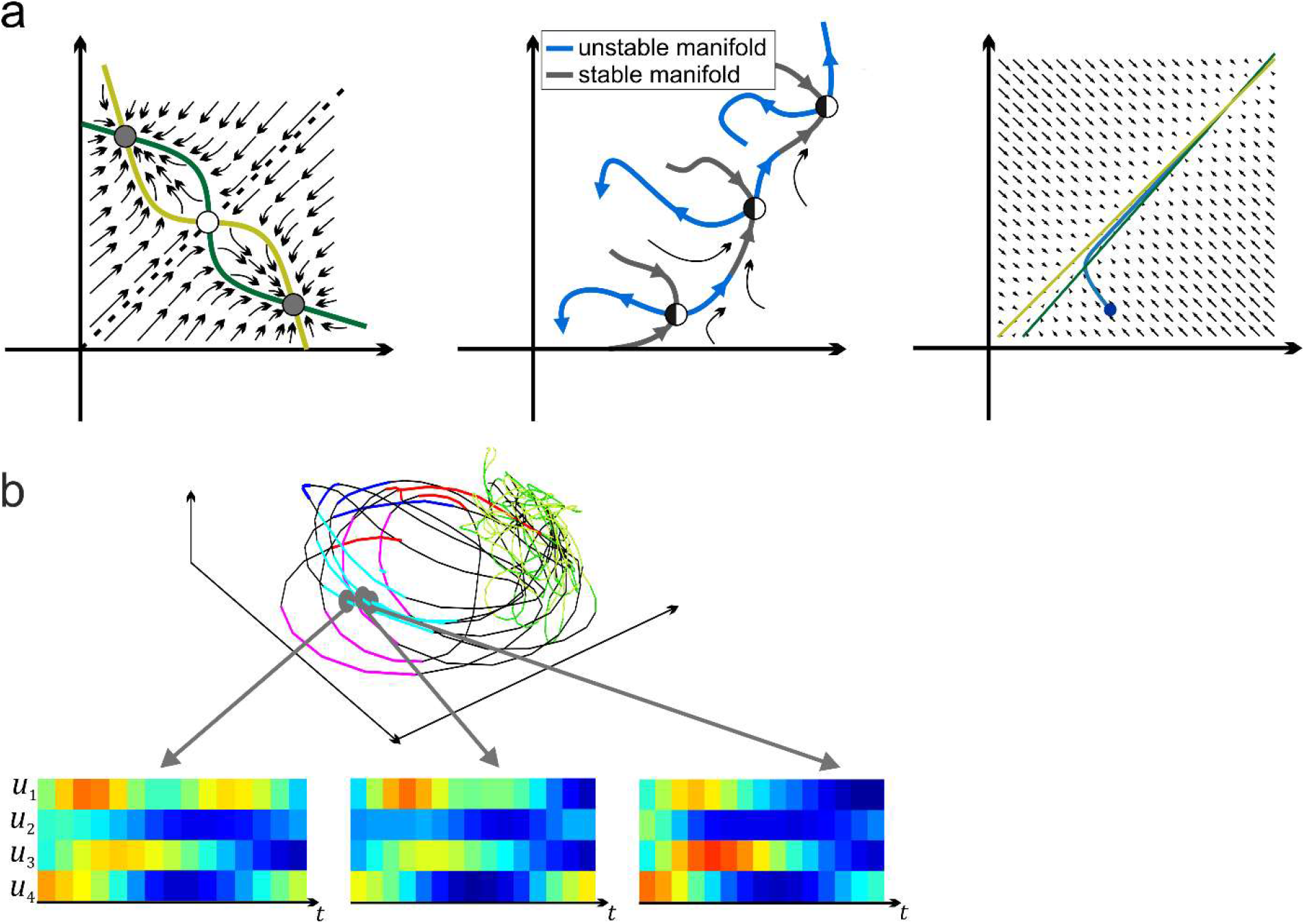
Interpretation of trained RNNs: Computational dynamics. (a) Different dynamical scenarios for working memory: Working memory may be implemented through multi-stability (left), separatrix cycles (center; cf. Rabinovich et al., 2008), or close-to-bifurcation dynamics in the vicinity of a detuned line attractor (right, adapted from Schmidt et al., 2021). An RNN trained on physiological data may be able to differentiate between these possibilities. (b) Dynamical phenomena map onto physiological processes: For instance, nearby trajectories in a neural “state space” correspond to similar spatio-temporal activity patterns (assemblies). Based on MSU data from the anterior cingulate cortex from rats performing a radial arm maze task; data kindly provided by Dr. Chris Lapish (Indianapolis Univ.).

Fig. 10 illustrates some of the possibilities for endowing the model with physiological and/or anatomical interpretability. Commonly, we couple a latent RNN as in eq. 1 to the actually measured time series through an observation (or decoder) model (see also Fig. 6): For instance, in the simplest case by a linear Gaussian model of the form

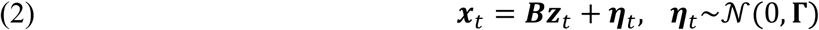

where ***x***_t_ are the multivariate observation (e.g., convolved spike trains), ***z***_t_ are the RNN’s latent states (cf. eq. 1), and ***B*** is a *factor loading matrix*. If the decoder model we use to link the latent RNN to the measured data takes the form of a generalized linear model (GLM), then we can directly interpret the entries in the GLM’s factor loading matrix ***B*** as in conventional (Durstewitz, 2017b) or Gaussian-process (Yu et al., 2009) factor analysis: They reflect the joint loading of different observables (like recorded units and/or behavioral responses) on the same (nonlinear) latent RNN factors. For instance, in the example in Fig. 10a, the sets of units appearing reddish within one column of ***B*** would be active within the same cell assembly, with different columns assigned to different latent states and hence assemblies. If measurements were taken from different data modalities simultaneously (Kramer et al. 2022), the factor loading matrix would also allow to identify relations between these modalities; for instance, between cell assemblies and behavioral variables.

An interesting option is to restrict the structure of ***B***: We could constrain ***B*** such that only subsets of RNN latent states are allowed to map to specified subsets of observations. Through this we could assign particular *semantic roles* to defined subsets of latent states: Suppose we have recordings from different brain areas or cortical layers and we constrain ***B*** such that only latent states #1-10 connect to brain area A and states #11-20 to brain area B (Fig. 10b). In that case, the entries in the RNN’s connectivity matrix ***W*** would automatically assume interpretations in terms of intra- and inter-areal connection strengths, as shown in Fig. 10b. As another example, assume we were able to sort the recorded units into different classes like pyramidal cells and interneurons. By assigning subsets of latent states to only one or the other class and enforcing their outgoing weights to be only positive or negative, matrix ***W*** would yield information about excitatory and inhibitory microcircuits and connectivity motives (Fig. 10c). Algorithmically, such constraints are often straightforward and easy to implement. We may also choose to make our factor loading matrices ***B***_k_ *trial-specific* (i.e., assign a different ***B***_k_ to each (set of) trial/s *k*) while keeping the latent model the same across trials or recording batches. For instance, when we assume that the set (and number) of recorded units differs across trials or recording sessions (or animals), while the latent dynamics remains the same.

Yet, our main incentive in using RNNs lies in the detailed analysis of the DS phenomena behind any task-related neural computations. An illustration of what this could look like is given in Fig. 11a: Shown are three state spaces with vector fields that exemplify different hypotheses about the neuro-computational realization of working memory. Fig. 11a (left) represents the classical ‘multi-stability hypothesis’ of working memory, where different attractor states correspond to different memory items (Amit & Brunel, 1997; Durstewitz et al., 2000; Wang, 1999). Fig. 11a (center) implements the idea that working memory is supported by a separatrix cycle (also called heteroclinic channel) that gives rise to a sequence of semi-stable states which encode memory contents (Afraimovich et al., 2004; Balaguer-Ballester et al., 2011; Lapish et al., 2015; Rabinovich et al., 2008; Rajan et al., 2016). Fig. 11a (right), finally, is based on the hypothesis that working memory does not really require stable states (attractors), but only relatively slow and adaptable time constants that arise in the vicinity of a bifurcation (Nakahara & Doya, 1998). There is some experimental support for all three of these ideas, e.g. the classical observation of persistent activity in the first case (Funahashi, 1989; Fuster, 1973), sequential activity in prefrontal cortex and other areas in the second (Baeg et al., 2003; Harvey et al., 2012; Rajan et al., 2016), and slowly ramping (climbing) activity in the third (Quintana & Fuster, 1999; Rainer et al., 1999). The main point here is that we now could use RNNs to provide more direct evidence for each of these scenarios by examining their state spaces, vector fields, equilibrium points, and so on, after they have been trained to faithfully reproduce neurophysiological data. By systematically analyzing the system dynamics we may even uncover unexpected dynamical mechanisms we have not previously thought of. Through the observation model’s loading matrix it is then possible to link up the unraveling latent dynamics to the associated physiological processes, e.g. to cell assemblies, as illustrated in Fig. 11b.

The success of this endeavor partly hinges on the dynamical accessibility and interpretability of the RNN model. If the mathematical setup of the RNN employed is itself rather complex, as for instance in LSTMs, we need to revert to approximate numerical methods to find dynamical objects and structure of interest, like fixed points (Sussillo & Barak, 2013). Some of these (like unstable cycles) are non-trivial to locate by pure numerical techniques (Krauskopf et al., 2007). Specific RNN formulations (Brenner et al., 2022; Durstewitz, 2017a; Koppe et al., 2019), regularization criteria (Duncker et al., 2019; Schmidt et al., 2021), or joint-model training methods (Smith et al., 2021) have therefore been introduced to facilitate dynamical analysis. PLRNNs as introduced above (Durstewitz, 2017a; Koppe et al., 2019; Kramer et al., 2022; Schmidt et al., 2021) have a mathematical structure that makes them more amenable to rigorous DST analysis to begin with, without compromising expressiveness or performance (Brenner et al., 2022). For instance, fixed points and even cycles (stable, unstable, or half-stable) can be computed *exactly* and in part analytically for PLRNNs (Brenner et al., 2022; Schmidt et al., 2021). There are also rules for converting between equivalent discrete and continuous-time PLRNNs, which enables to harvest training algorithms and analysis tools from both domains (Monfared & Durstewitz, 2020a). More generally, PLRNNs belong to the class of piecewise-linear (PWL) maps which have been extensively studied in DST (Avrutin et al., 2019; Monfared & Durstewitz, 2020b; Patra, 2018). In the field of DS reconstruction, the particular challenge thus is to construct simple and mathematically tractable, yet expressive model architectures. This puts a larger burden on the design of efficient training algorithms which need to be able to cope with smaller and simpler models yet produce faithful reconstructions of the dynamics (Mikhaeil et al., 2022).

## 4. Outlook and future challenges

The field of model-based DS reconstruction is still in its infancy. So far we know only little about the empirical and theoretical conditions under which we can trust our model training algorithms to yield good approximations to the underlying DS. Many algorithms for DS reconstruction have been tested so far only on rather small benchmark models (like the 3d Lorenz equations), with little process and observation noise, and assuming access to all system variables (but see Barkarji et al. 2022; Brenner et al., 2022; Li et al., 2021; Mikhaeil et al., 2022). In reality, especially in neuroscience, we may be dealing with rather high levels of both intrinsic and observation noise. Systematic theoretical studies on how much noise any reconstruction algorithm can tolerate are therefore needed. This is particularly true if the observed data sets are rather small, as in fMRI or many behavioral studies where we are dealing with comparatively short time series, on the order of hundreds (reconstruction with such small time series has indeed been tested (Koppe et al., 2019; Kramer et al., 2022), but most commonly hundreds of thousands of time points are assumed to be available). In neuroscience we are also often dealing with non-continuous, non-Gaussian measurements, in contrast to the large majority of DS reconstruction algorithms that expect continuous Gaussian-like data (but see Kim et al., 2021; Kramer et al., 2022). For instance, spike trains are (potentially Poisson) point processes, and behavioral variables are often categorical. It is as yet unclear how much detailed information about the underlying dynamics we can, even in principle, retrieve from such types of observations (Kramer et al., 2022). A related question is by how much different types of data pre-processing (e.g., various filtering operations, kernel density smoothing of spike trains etc.) diminish or enhance our ability to reconstruct the true underlying DS.

Moreover, neural systems are extremely high-dimensional. Modern recordings techniques now routinely provide hundreds (MSU) to thousands (Ca^2+^ imaging or fMRI) of simultaneous time series observations (Machado et al., 2022; Paulk et al., 2022; Urai et al., 2022; Vogt, 2019). Even though it has been speculated that the relevant neural dynamics may be confined to much lower-dimensional manifolds (Altan et al., 2021; Jazayeri & Ostojic, 2021), the number of recorded cells often still remains a minuscule fraction of all dynamical variables we have in the biological substrate (e.g., alone the billions of neurons in a rodent brain, not to mention all the cellular and molecular processes). Hence, even with such large capacity measurements, and notwithstanding the ‘low-*d*-manifold’ hypothesis, it is still unclear whether any experiment has probed all relevant degrees of freedom that determine the dynamics. Hence, we may still deal with the problem of *partial observations* (Bakarji et al. 2022). Likewise, it is not quite clear whether phenomena like multi-stability (Durstewitz et al., 2000; Wang, 2001) or chaotic itinerancy (Tsuda, 2015) that have been hypothesized to play a big role in neural dynamics are easily consistent with the ‘low-*d*-manifold’ hypothesis. If it turns out we indeed need rather high-dimensional models for a faithful reconstruction of dynamics, this in turn brings on severe challenges for model analysis and visualization. A related question is how to extract from trained models statistics most informative about the dynamics (e.g., Lyapunov spectra, number of fixed points and cycles, properties of attractor basins etc.) that could be used in large group comparisons, e.g. different patient cohorts (Thome et al., 2021).

So far mostly very simple dynamical phenomena like fixed points (Kim et al., 2021), limit cycles (Russo et al., 2018) or line attractors (Mante et al., 2013) have been studied in neuroscience. This may be related to the fact that most common behavioral tasks engaged in neuroscience are quite reduced and simple (e.g., involving only two cues and response options). Although perhaps useful to isolate cognitive components, this seems far from the complexity and variety of the natural and social world, which require a high degree of behavioral flexibility, planning, problem solving, and forecasting abilities. It appears rather unlikely that the cognitive processes encountered in such more natural contexts could be reduced to hopping just between a few fixed point attractors. For instance, the biophysical, morphological, and synaptic heterogeneity, and the many strong nonlinearities present in the nervous system, easily give rise to chaotic activity (Durstewitz & Gabriel, 2007; Kramer et al., 2022; Landau & Sompolinsky, 2018; London et al., 2010; van Vreeswijk & Sompolinsky, 1996), at first glance incompatible with the simple fixed point picture (Pereira-Obilinovic et al., 2021) (as a note on the side, however, chaos at the level of spiking activity may not be incompatible with fixed points in some averaged quantity like firing rates). Indeed, using RNNs trained to reconstruct DS from MSU recordings we often observed that task-relevant activity only corresponded to transients of the dynamics, while asymptotically neural activity would converge to chaotic attractors (Fig. 9). Thus, more complex DST concepts may need to be utilized and explored as neuroscience continues to delve into more complex behaviors and natural environments.

Another fundamental challenge to many reconstruction algorithms we would like to mention is that neuroscience data are often highly non-stationary, involving systematic trends and drifts (Clopath et al., 2017; Ecker et al., 2010), long time scale oscillations (Buzsáki, 2006; Buzsaki & Draguhn, 2004), slow changes in bodily, motivational or emotional states (Eldar & Niv, 2015; Mai et al., 2012), learning phenomena (Durstewitz et al., 2010; Russo et al., 2021), or slow changes in environmental inputs. There are various ways to deal with such non-stationarities in time series models, e.g. by treating the model parameters themselves as dynamical variables that can fluctuate across time (Park et al., 2015; Shimazaki et al., 2012; Zhao & Park 2020). However, as far as DS reconstruction is concerned, this topic has not been much explored yet. It is as yet unclear whether and how DS can be retrieved from time series measurements which entail slow parameter drifts (Patel & Ott, 2022). The problem here is partly related to the exploding/vanishing gradient issue briefly raised in sect. 2.3: For underlying chaotic systems, time series batches cannot be too long if divergence of trajectories and error gradients is to be avoided (Mikhaeil et al., 2022). Yet, this may hamper the detection of phenomena that evolve on slower time scales, as these may not be sufficiently represented in the short time batches used in training.

Finally, there is the issue of *generalization*: If our dynamic model is correct, it should be able to generalize *beyond* the data domain seen in training, sometimes labeled ‘out-of-distribution’ generalization in machine learning (in contrast to ‘mere’ out-of-sample prediction). The problem of drifting parameters considered above may be seen as one facet of this issue, as a good model should be able to predict dynamics beyond the bifurcation point within unseen parameter regimes (Patel & Ott, 2022). Experimental manipulations could be designed to test such out-of-distribution predictions: For instance, if we have a representation of different neuron types or cortical layers in our model (as in Fig. 10), then optogenetic silencing or activation of the respective cell populations could be simulated in the RNN. Another facet of out-of-distribution generalization occurs in *multi-stable* systems: If an RNN has seen only data from a subset of the system’s attractor basins, could it still generalize to those basins and dynamical regimes not observed during training? If the training delivered a proper DS reconstruction, this should be the case, but in practice likely more work on topological theory for RNN-based DS reconstruction and on training algorithms will be necessary.

In summary, there are quite a number of problems that still need to be addressed and researched in the area of model-based DS reconstruction in general, and neuroscience in particular. Many of these are likely to require close interaction between experimentalists and theoreticians as experiments may need to be purpose-designed to tackle some of these outstanding issues.

## Acknowledgements

This work was funded by the German Research Foundation (DFG) through individual grants Du 354/10-1 and Du 354/15-1 to DD, and Du 354/14-1 within the DFG research cluster FOR-5159 (“Resolving prefrontal flexibility”). We thank Jonas Mikhaeil for providing detailed feedback and suggestions on this article, and Lukas Judith for providing the EEG simulations.

